# An Efficient, Scalable and Exact Representation of High-Dimensional Color Information Enabled via de Bruijn Graph Search

**DOI:** 10.1101/464222

**Authors:** Fatemeh Almodaresi, Prashant Pandey, Michael Ferdman, Rob Johnson, Rob Patro

## Abstract

The colored de Bruijn graph (cdbg) and its variants have become an important combinatorial structure used in numerous areas in genomics, such as population-level variation detection in metagenomic samples, large scale sequence search, and cdbg-based reference sequence indices. As samples or genomes are added to the cdbg, the color information comes to dominate the space required to represent this data structure.

In this paper, we show how to represent the color information efficiently by adopting a hierarchical encoding that exploits correlations among color classes — patterns of color occurrence — present in the de Bruijn graph (dbg). A major challenge in deriving an efficient encoding of the color information that takes advantage of such correlations is determining which color classes are close to each other in the high-dimensional space of possible color patterns. We demonstrate that the dbg itself can be used as an efficient mechanism to search for approximate nearest neighbors in this space. While our approach reduces the encoding size of the color information even for relatively small cdbgs (hundreds of experiments), the gains are particularly consequential as the number of potential colors (i.e. samples or references) grows to thousands of experiments.

We apply this encoding in the context of two different applications; the implicit cdbg used for a large-scale sequence search index, Mantis, as well as the encoding of color information used in population-level variation detection by tools such as Vari and Rainbowfish. Our results show significant improvements in the overall size and scalability of representation of the color information. In our experiment on 10,000 samples, we achieved more than 11*×* better compression compared to RRR.

## 1 Introduction

The colored de Bruijn graph (cdbg) [1], an extension of the classical de Bruijn graph [2–4], is a key component of a growing number of genomics tools. Augmenting the traditional de Bruijn graph with “color” information provides a mechanism to associate meta-data, such as the raw sample or reference of origin, with each *k*-mer. Coloring the de Bruijn graph enables it to be used in a wide range of applications, such as large-scale sequence search [5–9], though some [6–8] do not explicitly couch their representations in the language of the cdbg), population-level variation detection [10–12], traversal and search in a pan-genome [13], and sequence alignment [14]. The popularity and applicability of the cdbg has spurred research into developing space-efficient and high-performance data-structure implementations.

An efficient and fast representation of cdbg requires optimizing both the de Bruijn graph and the color information. While there exist efficient and scalable methods for representing the topology of the de Bruijn graph [4, 15–19] with fast query time, a scalable and exact representation of the color information has remained a challenge. Recently, Mustafa et al. [20] has tackled this challenge by relaxing the exactness constraints — allowing the returned color set for a *k*-mer to contain extra samples with some controlled probability — but it is not immediately clear how this method can be made exact.

Specifically, existing exact color representations suffer from large sizes and a fast growth rate that leads them to dominate the total representation size of the cdbg with even a moderate number of input samples(see Figure 3b). As a result, the color information grows to dominate the space used by all these indexes and limits their ability to scale to large input data sets.

Iqbal et al. introduced cdbgs [1] and proposed a hash-based representation of the de Bruijn graph in which each *k*-mer is additionally tagged with the list of reference genomes in which it is contained. Muggli et al. reduced the size of the cdbg in VARI [10] by replacing the hash map with BOSS [17] (a BWT-based [21] encoding of the de Bruijn graph that assigns a unique ID to each *k*-mer) and using a boolean matrix indexed by the unique *k*-mer ID and genome reference ID to indicate occurrence. They reduced the size of the occurrence matrix by applying off-the-shelf compression techniques RRR [22] and Elias-Fano [23] encoding. Rainbowfish [12] shrank the color table further by ensuring that rows of the color matrix are unique, mapping all *k*-mers with the same color information to a single row, and assigning row indices based on the frequency of each occurrence pattern. However, despite these improvements, the scalability of the resulting structure remains limited because even after eliminating redundant colors, the space for the color table grows quickly to dominate the total space used by these data structures.

We observe that, in real biological data, even when the number of distinct color classes is large, many of them will be near each other in terms of the set of samples or references they encode. That is, the color classes tend to be highly correlated rather than uniformly spread across the space of possible colors. There are intuitive reasons for such characteristics. For example, we observe that adjacent *k*-mers in the de Bruijn graph are extremely likely to have either identical or similar color classes, enabling storage of small deltas instead of the complete color classes. This is because *k*-mers adjacent in the de Bruijn graph are likely to be adjacent (and hence present) in a similar set of input samples. In the context of sequence-search, because genomes and transcriptomes are largely preserved across organs, individuals, and even across related species, we expect two *k*-mers that occur together in one sample to be highly correlated in their occurrence across many samples. Thus, we can take advantage of this correlation when devising an efficient encoding scheme for the cdbg’s associated color information.

In this paper, we develop a general scheme for efficient and scalable encoding of the color information in the cdbg by encoding color classes (i.e. the patterns of occurrence of a *k*-mer in samples) in terms of their differences (which are small) with respect to some “neighboring” color class. The key technical challenge, solved by our work, is efficiently searching for the neighbors of color classes in the high-dimensional space of colors by leveraging the observation that similar color classes tend to be topologically close in the underlying de Bruijn graph. We construct a weighted graph on the color classes in the cdbg, where the weight of each edge corresponds to the space required to store the delta between its endpoints. Finding the minimum spanning tree (MST) of this graph gives a minimal delta-based representation. Although reconstructing a color class on this representation requires a walk to the MST root node, abundant temporal locality on the lookups allows us to use a small cache to mitigate the performance impact, yielding query throughput that is essentially the same as when all color classes are represented explicitly.

We showcase the generality and applicability of our color class table compression technique by demonstrating it in two computational biology applications: sequence search and variation detection. We compare our novel color class table representation with the representation used in Mantis [5], a state-of-the-art large-scale sequence-search tool that uses a cdbg to index a set of sequencing samples, and the representation used in Rainbowfish [12], a state-of-the-art index to facilitate variation detection over a set of genomes. We show that our approach maintains the same query performance while achieving over 11*×* and 2.5*×* storage savings relative to the representation previously used by these tools.

## 2 Methods

This section describes our compact cdbg representation. We first define cdbgs and briefly describe existing compact cdbg representations. We then outline the high-level idea behind our compact representation and explain how we use the de Bruijn graph to efficiently build our compact representation. Finally, we describe implementation details and optimizations to our query algorithm.

### 2.1 Colored de Bruijn graphs

De Bruijn graphs are widely used to represent the topological structure of a set of *k*-mers [2, 19, 24–29]. The de Bruijn graph induced by a set of *k*-mers is defined below.

#### Definition 1.

*Given a set E of k-mers, the de Bruijn graph induced by E has edge set E, where each k-mer (or edge) connects its two* (*k−*1)*-length substrings (or vertices).*

Cdbgs extend the de Bruijn graph by assigning a *color class C*(*x*) to each edge (or node) *x* of the de Bruijn graph. The color class *C*(*x*) is a set drawn from some universe *U*. Examples of *U* and *C*(*x*) are

– Sometimes, *U* is a set of reference genomes, and *C*(*x*) is the subset of reference genomes containing *k*-mer *x* [10, 12, 14, 30].
– Sometimes, *U* is a set of *reads*, and *C*(*x*) is the subset of reads containing *x* [31–33].
– Sometimes, *U* is a set of sequencing experiments, and *C*(*x*) is the subset of sequencing experiments containing *x* [5–8].

The goal of a cdbg representation is to store *E* and *C* as compactly as possible^3^, while supporting the following operations efficiently:

– *Point query.* Given a *k*-mer *x*, determine whether *x* is in *E*.
– *Color query.* Given a *k*-mer *x∉E*, return *C*(*x*).

Given that we can perform point queries, we can traverse the de Bruijn graph by simply querying for the 8 possible predecessor/successor edges of an edge. This enables us to implement more advanced algorithms, such as bubble calling [34].

Many cdbg representations typically decompose, at least logically, into two structures: one structure storing a de Bruijn graph and associating an ID with each *k*-mer, and one structure mapping these IDs to the actual color class [10, 12, 35]. The individual color classes can be represented as bit-vectors, lists, or via a hybrid scheme [36]. This information is typically compressed [22, 37, 38].

Our paper follows this standard approach, and focuses exclusively on reducing the space required for the structure storing the color information. We propose a new representation that, given a color ID, can return the corresponding color efficiently. Although, as a matter of convenience, we pair our color table representation with the de Bruijn graph structure representation of the counting quotient filter [35] as used in Mantis [5], our proposed color table representation can be paired with other de Bruijn graph representations.

### 2.2 A similarity-based cdbg representation

The key observation behind our compressed color-class representation is that the color classes of *k*-mers that are adjacent in the de Bruijn graph are likely to be very similar. Thus, rather than storing each color class explicitly, we can store only a few color classes explicitly and, for all the remaining color classes, we store only their differences from other color classes. Because the differences are small, the total space used by the representation will be small.

Motivated by the observation above, we propose to find an encoding of the color classes that takes advantage of the fact that most color classes can be represented in terms of only a small number of edits (i.e., flipping the parity of only a few bits) with respect to some neighbor in the high-dimensional space of the color classes. In fact, this idea was explored by Bookstein and Klein [39] in the context of information retrieval, where an elegant solution was proposed. We describe this approach below, in the context of the set of correlated bit vectors (i.e. color classes) that we wish to encode.

We construct our compressed color class representation as follows (see Figure 1). For each edge *x* of the de Bruijn graph, let *C*(*x*) be the color class of *x*. Let *C* be the set of all color classes ({*C*(*x*)*| x∈* dbg}) that occur in the de Bruijn graph. We first construct a graph with vertex set *C* and edge set reflecting the adjacency relationship implied by the de Bruijn graph. In other words, there is an edge between color classes *c*_1_ and *c*_2_ if there exist adjacent edges (i.e. incident on the same node) *x* and *y* in the de Bruijn graph such that *c*_1_ =*C*(*x*) and *c*_2_ =*C*(*y*). These edges indicate color classes that are likely to be similar, based on the structure of the de Bruijn graph. We then add a special node ∅ to the color class graph, which is connected to every node. We set the weight of every edge in the color class graph to be the Hamming distance between its two endpoints (where we view color classes as bit vectors and ∅ is the all-zeros bit vector).

**Fig. 1:**
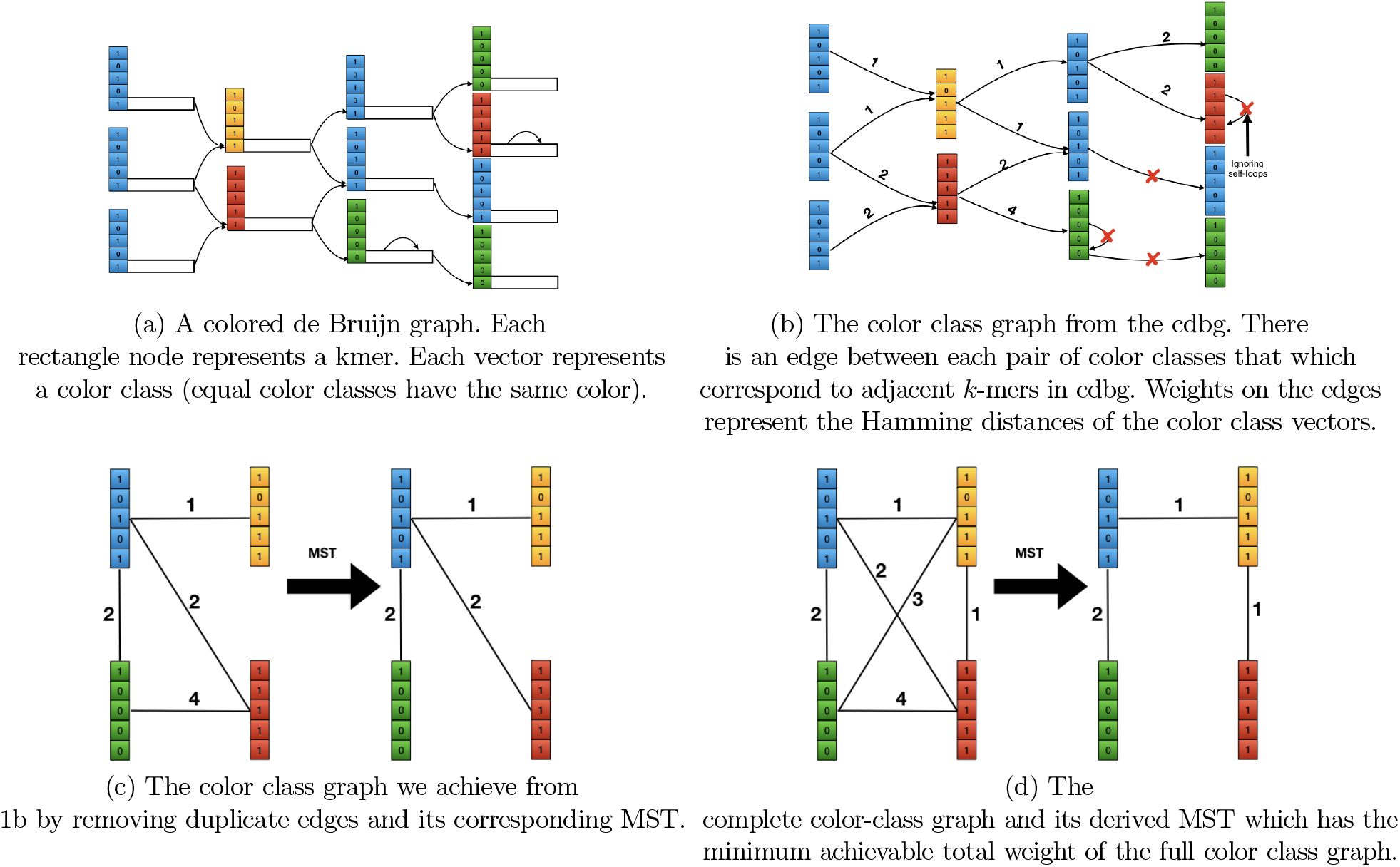
Encoding color classes by finding the MST of the color class graph derived from cdbg. The order of the process is 1a, 1b, and 1c and the optimal achievable MST is shown in 1d for comparison. Since we never observe the edge between any *k*-mers from color classes green and yellow in cdbg, we won’t have the edge between color classes green and yellow and therefore, our final MST is not equal to the best MST we can get from a complete color class graph.

We then compute a minimum spanning tree of the color class graph and root the tree at the special ∅ node. Note that, because the ∅ node is connected to every other node in the graph, the graph is connected and hence an MST is guaranteed to exist. By using a minimum spanning tree, we minimize the total size of the differences that we need to store in our compressed representation.

We then store the MST as a table mapping each color class ID to the ID of its parent in the MST, along with a list of the differences between the color class and its parent. For convenience we can view the list of differences between color class *c*_1_ and color class *c*_2_ as a bit vector *c*_1_ ⊕ *c*_2_, where ⊕ is the bit-wise exclusive-or operation. To reconstruct a color class given its ID *i*, we simply xor all the difference vectors we encounter while walking from *i* to the root of the MST.

### 2.3 Implementation using succinct data structures

Assuming we have *|C|* color classes, *|U|* colors, and an MST with total weight of *w* over the color class graph, we store all the information required to retrieve the original color bit-vector for each color class ID based on the MST structure into three data structures:

– **Parent vector**: This vector contains *|C|* slots, each of size ⌈log2 *C*⌉. The value stored in index *I* represents the parent color class ID of the color class with index *i* in the MST.
– **Delta vector**: This vector contains *w* slots, each of size ⌈log2*|U|*⌉. For each pair of parent and child in the parent vector, we compute a vector of the indices at which they differ. The delta vector is the concatenation of these per-edge delta vectors, ordered by the ID of the source of the edge. Note that the per-edge delta vectors will not all be of the same length, because some edges have larger weight than others. Thus, we need an auxiliary data structure to record the boundaries between the per-edge deltas within the overall delta vector.
– **Boundary bit-vector**: This vector contains *w* bits, where a set bit indicates the boundary between two delta sets within the delta vector. To find the starting position, within the delta vector, of the per-edge delta list for the MST edge with source ID *i*, we perform *select(i)* on the boundary vector. Select returns the position of the *i*th one in the boundary vector.

#### Query of the MST-based representation

Figure 2 shows how queries proceed using this encoding. We start with an empty accumulator bit vector and a color class ID *i* for which we want to compute the corresponding color class. We perform a select query for *i* and *i*+1 in the boundary bit-vector to get the boundaries of *i*’s difference list in the delta vector. We then iterate over its difference list and flip the indicated bits in our accumulator. We then set *i←*PARENT[*i*] and repeat until *i* becomes 0, which indicates that we have reached the root. At this point, the accumulator will be equal to the bit vector for color class *i*.

**Fig. 2:**
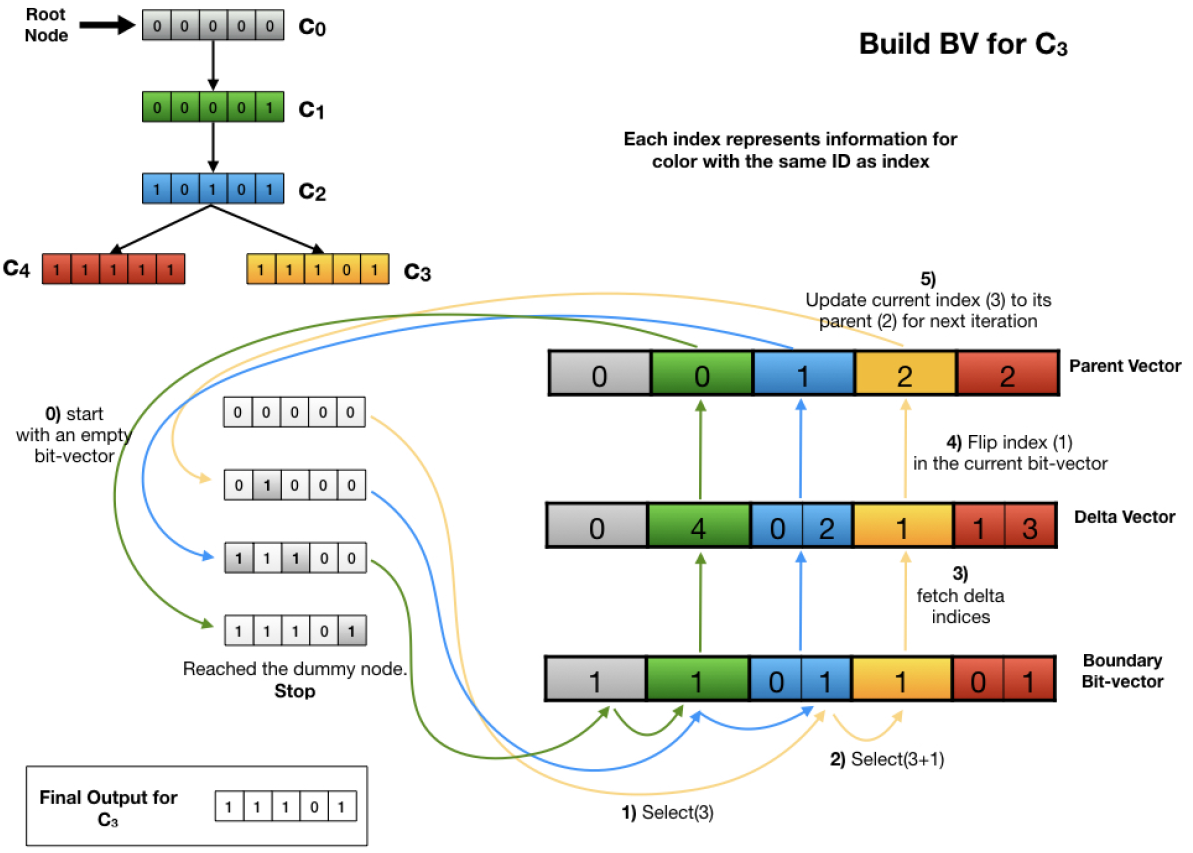
The conceptual MST (top-left), the data structure to store the color information in the format of an MST (right). This figure also illustrates the steps required to build one of the color vectors (*C*_3_) at the leaf of the tree. Note that the query process shown here does not depict the caching policy we apply in practice.

### 2.4 Integration in Mantis

Once constructed, our MST-based color class representation is a drop-in replacement for the current color class representations used in several existing tools, including Mantis [5] and Rainbowfish [12]. Their existing color class tables support a single operation—querying for a color class by its ID—and our MST-based representation supports exactly the same operation.

For this paper, we integrated our MST-based representation into Mantis. The same space savings can be achieved in other tools, particularly Rainbowfish, which has a similar color-class encoding as Mantis.

#### Construction

We construct our MST-based color-class representation as follows. First, we run Mantis to build its default representation of the cdbg. We then build the color-class graph by walking the de Bruijn graph and adding all the corresponding edges to the color-class graph. The edge set is typically much smaller than the de Bruijn graph (because many de Bruijn graph edges may map to the same edge in the color-class graph), so this can be done in RAM. Note that we do not compute the weights of the edges during this pass, because that would require having all of the large color-class bit vectors in memory in order to compute their Hamming distance.

In the second pass, we traverse the edge set and compute the weight of each edge. To minimize RAM usage during this phase, we sort the edges and iterate over them in a “blocked” fashion. Specifically, Mantis stores the color class bit vectors on-disk sequentially by ID, grouped into blocks of roughly 6GBs each. We sort the edges lexicographically by their source and destination block. We then load all pairs of blocks and compute the weights of all the edges between the two blocks currently in memory. At all times, we need only two blocks of color class vectors in memory. Given the weighted graph, we compute the MST and make one final pass to determine the relevant delta lists and encode our final MST structure.

#### Parallelization

We note that, after having constructed the Mantis representation, most phases of the the MST construction algorithm are trivially parallelized. MST construction decomposes into three phases: (1) color-class graph construction, (2) MST computation, and (3) color-class representation generation. We parallelize graph construction and color-class representation generation. The MST computation itself is not parallelized.

We parallelized the determination of edges in the color-class graph by assigning each thread a range of the k*−mer*-to-color-class-ID map. Each thread explores the neighbors of the *k*-mers that appear in its assigned range, and any redundant edges are deduplicated when all threads are finished. Similarly, we parallelized the computation of edge weights and the extraction of the delta vectors that correspond to each edge in the MST. Given the list of edges sorted lexicographically by their endpoints (determined during the first phase), it is straightforward to partition the work for processing batches of edges across many threads. It is possible, of course, that the batches will display different workloads and that some threads will complete their assigned work before others. We have not yet made any attempt to optimize the parallel construction of the MST in this regard, though many such optimizations are likely possible.

#### Accelerating queries with caching

The encoded MST is not a balanced tree, so decoding a color bit-vector might require walking a long path to the root, which negatively impacts the query time. Additionally, considering the fact that the frequency distribution of the color classes is very skewed, some of the color classes are more popular or have more children and, therefore, are in the path of many more nodes. We take advantage of these data characteristics by caching the most recent queried color bit-vectors. Every time we walk up the tree, if the color bit-vector for a node is already in the cache, our query algorithm stops at that point and applies all the deltas to this bit-vector instead of the zero bit-vector of the root. This caching approach significantly improves the query time, resulting in the final query time required to decode a color class being marginally faster than direct RRR access.

The cache policy is designed with the tree structure of our color-class representation in mind. Specifically, we want to cache nodes near the leaves, but not so close to the leaves that we end up caching essentially the entire tree. Also, we don’t want to cache infrequently queried nodes. Thus we use the following caching policy: all queried nodes are cached. Furthermore, we cache interior nodes visited during a query as follows. If a query visits a node that has been visited by more than 10 other queries and is more than 10 hops away from the currently queried item, then we add that node to the cache. If a query visits more than one such node, we add the first one encountered.

In our experiments, we used a cache of 10,000 nodes and managed the cache using a FIFO policy.

### 2.5 Comparison with brute-force and approximate-nearest-neighbor-based approaches

Our MST-based color-class representation uses the de Bruijn graph as a hint as to which color classes are likely to be similar. This leads to the natural question: how good are the hints provided by the de Bruijn graph?

One could imagine alternatively constructing the MST on the complete color-class graph. This would yield the absolutely lowest-weight spanning tree on the color classes. Unforunately, no MST algorithm runs in less than Ω(*|E|*) time, so this would make our construction time quadratic in the number of color classes. The number of color classes in our experiments range from 10^6^ to 10^9^, so the number of edges in the complete color-class graph would be on the order of 10^12^ to 10^18^, or possibly even more, making this algorithm impractical for the largest data sets considered in this paper.

Alternatively, we could try to use an approximate nearest-neighbor algorithm to find pairs of color classes with small Hamming distance. As an experiment, we implemented an approximate nearest neighbor algorithm that bucketed color classes by their projection into a smaller-dimensional subspace. Nearest-neighbor queries were computed by searching within the queried item’s bucket. Results were disappointing. Even on small data sets, the average distance between the queried item and the returned neighbor was several times larger than the average distance found using the neighbors suggested by the de Bruijn graph. Thus, we did not pursue this direction further.

## 3 Evaluation

In this section we evaluate our MST-based representation of the color information in the cdbg. All our experiments use Mantis with our integrated MST-based color-class representation.

**Evaluation Metrics** We evaluate our MST-based representation on the following parameters:

– **Scalability.** How does our MST-based color-class representation scale in terms of space with increasing number of input samples, and how does it compare to the existing representations of Mantis?
– **Construction time.** How long does it take – in addition to the original construction time for building cdbg– to build our MST-based color-class representation?
– **Query performance.** How long does it takes to query the cdbg using our MST-based color-class representation?

### 3.1 Experimental procedure

#### System Specifications

Mantis takes as input a collection of *squeakr* files [40]. Squeakr is a *k*-mer counter that takes as input a collection of fastq files and produces as output, a single file with a compact hash table mapping each *k*-mer to the number of times it occurs in the input files. As is standard in evaluations of large-scale sequence search indexes, we do not benchmark the time required to construct these filters.

The data input to the construction process was stored on 4-disk mirrors (8 disks total). Each is a Seagate 7200rpm 8TB disk (ST8000VN0022). They were formatted using ZFS and exported via NFS over a 10Gb link. We used different systems to run and evaluate time, memory, and disk requirements for the two steps of preprocessing and index building as was done by Pandey et al. [5].

For index building and query benchmarks, we ran all the experiments on the same system used in Mantis [5], an Intel(R) Xeon(R) CPU (E5-2699 v4 @2.20GHz with 44 cores and 56MB L3 cache) with 512GB RAM and a 4TB TOSHIBA MG03ACA4 ATA HDD running Ubuntu 16.10 (Linux kernel 4.8.0-59-generic). Constructing the main index was done using a single thread, and the MST construction was performed using 16 threads. Query benchmarks were also performed using a single thread.

#### Data to evaluate scalability and comparison to Mantis

We integrated and evaluated our MST-based color-class representation within Mantis, so we briefly review Mantis here. Mantis builds an index on a collection of unassembled raw sequencing data sets. Each data set is called a *sample*. The Mantis index enables fast queries of the form, “Which samples contain this *k*-mer,” and “Which samples are likely to contain this string of bases?” Mantis takes as input one *squeakr* file per sample [40]. A squeakr file is a compact hash table mapping each *k*-mer to the number of times it occurs within that sample.

Given the input files, Mantis constructs an index consisting of two files: a map from *k*-mer to color-class ID, and a map from color-class ID to the bit vector encoding that color class. The first map is stored as a *counting quotient filter* (CQF), which is the same compact hash table used by Squeakr. The color-class map is an RRR-compressed bit vector.

Recall that our construction process is implemented as a post-processing step on the standard Mantis color-class representation. For construction times, we report only this post-processing step. This is because our MST-based color-class representation is a generic tool that can be applied to many cdbg representations other than Mantis, so we want to isolate the time spent on MST construction.

To test the scalability of our new color class representation, we used a set of 10,000 human RNA-seq short-read experiments downloaded from European Nucleotide Archive(ENA) [41] in gzipped FASTQ format. This set includes the 2,586 sequencing samples from human blood, brain, and breast tissues (BBB) originally used by [42] and also used in the subsequent work [7, 8, 36], including Mantis [5]. The remaining 7, 414 experiments were selected randomly from the bulk human RNA-seq experiments downloaded from ENA. The full list of experimental identifiers can be obtained from https://github.com/COMBINE-lab/color-mst/blob/master/input_lists/shuffled_10k_paired. The total size of all experiments (gzipped) is 25.6TB.

In order to eliminate spurious *k*-mers that occur with insignificant abundance within a sample, the squeakr files are filtered to remove low-abundance *k*-mers. We adopted the same cutoff policy originally proposed by Solomon and Kingsford [42], by discarding *k*-mers that occur less than some threshold number of time. The thresholds are determined according to the size (in bytes) of the gzipped sample, and the thresholds are given in Table 1. We adopt a value of *k* =23 for all experiments.

**Table 1:** Minimum number of times a *k*-mer must appear in an experiment in order to be counted as abundantly represented in that experiment (taken from the SBT paper).

| Min size | Max size | Cutoff | # of experiments<br>with specified threshold |
| --- | --- | --- | --- |
| 0 | $\leq 300\text{MB}$ | 1 | 2,784 |
| $> 300\text{MB}$ | $\leq 500\text{MB}$ | 3 | 798 |
| $> 500\text{MB}$ | $\leq 1\text{GB}$ | 10 | 1,258 |
| $> 1\text{GB}$ | $\leq 3\text{GB}$ | 20 | 2,296 |
| $> 3\text{GB}$ | ∞ | 50 | 2,864 |

### 3.2 Evaluation results

#### Scalability of the new color class representation

Figure 3a and Table 2 show how the size of our MST-based color-class representation scales as we increase the number of samples indexed by Mantis. For comparison, we also give the size of Mantis’ RRR-compression-based color-class representation. Figure 3a also plots the size of the CQF that Mantis uses to map *k*-mers to color class IDs. We can draw several conclusions from this data:

– The MST-based representation is an order-of-magnitude smaller than the RRR-based representation.
– The gap between the RRR-based representation and the MST-based representation grows as we increase the number of input samples. This suggests that the MST-based representation grows asymptotically slower than the RRR-based representation.
– The MST-based color-class representation is, for large numbers of samples, about 5*×* smaller than the CQF. This means that representing the color classes is no longer the scaling bottleneck.

**Fig. 3:**
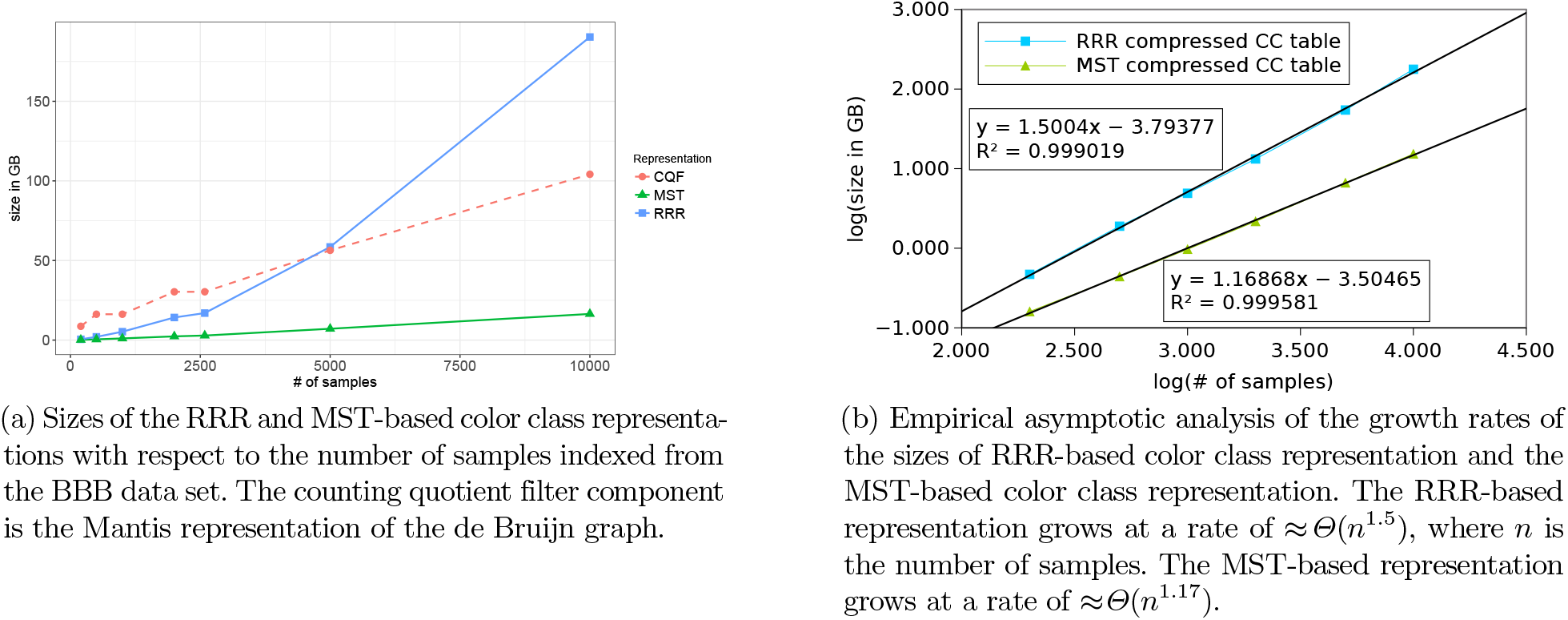
Size of the MST-based color-class representation vs. the RRR-based color-class representation.

**Table 2:**
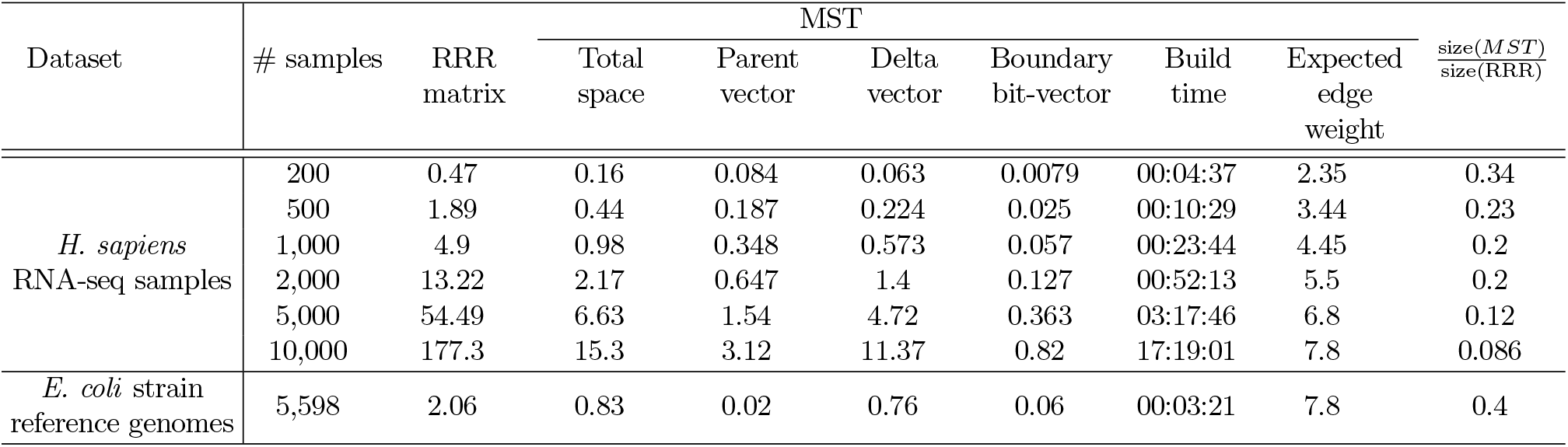
Space required for RRR and MST-based color class encodings over different numbers of samples (sizes in GB) and time required to build MST. Central columns break down the size of individual MST components.

Table 2 also shows the scaling rate of all elements of the MST representation, in addition to the ratio of MST over the color bit-vector. As expected, the list of deltas dominate the MST representation both in terms of total size and in terms of growth. Table 2 also shows the average edge weight of the edges in the MST. The edge weight grows approximately proportional to *Θ*(log(# of samples)) (i.e. every time we double the number of samples, the average edge weight increases by almost exactly 1). This suggests that our de Bruijn graph-based algorithm is able to find pairs of similar color classes.

To better understand the scaling of the different components of a cdbg representation, we plot the sizes of the RRR-based color-class representations and MST-based representations on a log-log scale in Figure 3b. Based on the data, the RRR-based representation appears to grow in size at a rate of roughly *Θ*(*n*^1.5^), whereas the new MST-based representation grows roughly at a rate of *Θ*(*n*^1.17^). This explains why the RRR-based representation grows to dwarf the CQF (which grows roughly linearly) and become the bottleneck to scaling to larger data sets, whereas the MST-based representation does not. With the MST-based representation, the CQF itself is now the bottleneck.

Finally, Figure 3a shows the size of the RRR-and MST-based color-class representations for the *E. coli* data set analyzed in the Rainbowfish paper. As the table shows, our MST-based color-class representation is able to effectively compress genomic color data in addition to RNA-seq color data.

#### Index Building Evaluation

The “Build time” column in Table 2 shows the time required to build our MST-based color-class representation from Mantis’ RRR-based representation. All builds used 16 threads.

Overall, the MST construction time is only a tiny fraction of the overall time required to build the Mantis index from raw fastq files. The vast bulk of the time is spent processing the fastq files to produce filtered squeakrs. This step was performed on a cluster of 150 machines over roughly one week. Thus MST construction represents less than 1% of the overall index build time.

#### Query Evaluation

We evaluate query speed in the following manner. We select random subsets, of increasing size, of transcripts from the human transcriptome, and query the Mantis index to determine the set of experiments containing each of these transcripts. Mantis answers transcript queries as follows. For each *k*-mer in the transcript, it computes the set of samples containing that *k*-mer. It then reports a sample as containing a transcript if the sample contains more than *Θ* fraction of the *k*-mers in the transcript, where *Θ* is a user-adjustable parameter. Note that, for Mantis, the *Θ* threshold is applied at the very end. Mantis first computes, for each sample, the fraction of *k*-mers that occur in that sample, and then filters as a last step. Thus the query times reported here are valid for any *Θ*.

Table 3 reports the query performance of both the RRR and MST-based Mantis indexes. Despite the vastly-reduced space occupied by the MST-based index, and the fact that the color class decoding procedure is more involved, query in the MST-based index is slightly faster than querying in the RRR-based index. The average query time in both RRR-based and MST-based index is 0.08 sec / query.

**Table 3:**
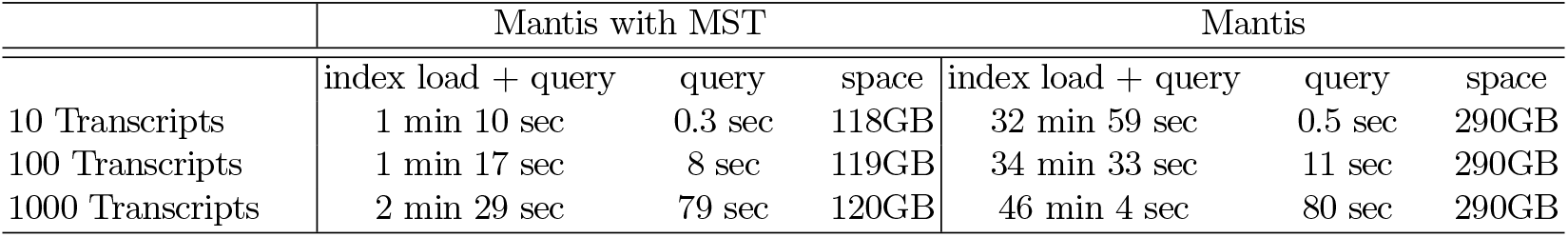
Query time and resident memory for mantis using the MST-based representation for color information and the original mantis (using RRR-compressed color classes) over 10,000 experiments. The “query” column provides just the time taken to execute all queries (as would be required if the index was already loaded in e.g. a server-based search tool). Note that, in resident memory usage for the MST-based representation, the counting quotient filter always dominates the total required memory.

|  | Mantis with MST |  |  | Mantis |  |  |
| --- | --- | --- | --- | --- | --- | --- |
|  | index load + query | query | space | index load + query | query | space |
| 10 Transcripts | 1 min 10 sec | 0.3 sec | 118GB | 32 min 59 sec | 0.5 sec | 290GB |
| 100 Transcripts | 1 min 17 sec | 8 sec | 119GB | 34 min 33 sec | 11 sec | 290GB |
| 1000 Transcripts | 2 min 29 sec | 79 sec | 120GB | 46 min 4 sec | 80 sec | 290GB |

Mantis queries are much faster than in other large-scale sequence search data structures, and our MST-based color-class representation doesn’t change that.

## 4 Discussion and Conclusion

We have introduced a novel exact representation of the color information associated with the cdbg. Our representation yields large improvements in terms of representation size when compared to previous state-of-the-art approaches. While our MST-based representation is much smaller, it still provides rapid query and can, for example, return the query results for a transcript across an index of 10,000 RNA-seq experiments in ~ 0.08 sec / query. Further, the size benefit of our proposed representation over that of previous approaches appears to grow with the number of color classes being encoded, meaning it is not only much smaller, but also much more scalable. Finally, the representation we propose is, essentially, a stand-alone encoding of the cdbg’s associated color information, making this representation conceptually easy to integrate with any tool or method that needs to store color information over a large de Bruijn graph.

Though it is not clear how much further the color information can be compressed while maintaining a lossless representation, this is an interesting theoretical question. It may be fruitful to approach this question from the perspective suggested by Yu et al. [43], of evaluating the metric entropy, fractal dimension and information-theoretic of the space of color classes. Practically, however, we have observed that, at least in our current system, Mantis, for large-scale sequence search, the counting quotient filter, which is used to store the topology of the de Bruijn graph and to associate color class labels with each *k*-mer, has become the new scalability bottleneck. Here, it may be possible to reduce the space required by this component by making use of some of the same observations we relied upon to allow efficient color class neighbor search. For example, because many adjacent *k*-mers in the de Bruijn graph share the same color class ID, it is likely possible to encode this label information sparsely across the de Bruijn graph, taking advantage of the coherence between topologically nearby *k*-mers. Further, to allow scalability to truly-massive datasets, it will likely be necessary to make the system hierarchical, or even to adopt a more space-efficient (and domain-specific) representation of the underlying de Bruijn graph. Nonetheless, because we have designed our color class representation as essentially orthogonal to the de Bruijn graph representation, we anticipate that we can easily integrate this approach with improved representations of the de Bruijn graph.

Mantis with the new MST-based color class encoding is written in `C++17` and is available at https://github.com/splatlab/mantis.

## Acknowledgments

This work was supported by the US National Science Foundation grants BIO-1564917, CCF-1439084, CCF-1452904, CCF-1716252, CCF-1750472, CNS-1408695, and CNS-1763680, and the US National Institutes of Health grant R01HG009937. The experiments were conducted with equipment purchased through NSF CISE Research Infrastructure Grant Number 1405641.

## Footnotes

3 The nodes of the de Bruijn graph are typical stored implicitly, because the node set is simply a function of *E*.

